# Fibroblast-like synoviocytes orchestrate daily rhythmic inflammation in arthritis

**DOI:** 10.1101/2024.04.02.587439

**Authors:** Polly Downton, Suzanna H. Dickson, David W. Ray, David A. Bechtold, Julie E. Gibbs

**Affiliations:** Centre for Biological Timing, Faculty of Biology, Medicine and Health, University of Manchester, Manchester, M13 9PT, UK; NIHR Oxford Health Biomedical Research Centre, and NIHR Oxford Biomedical Research Centre, John Radcliffe Hospital, Oxford, OX3 9DU, UK; Oxford Centre for Diabetes, Endocrinology and Metabolism, and Oxford Kavli Centre for Nanoscience Discovery, University of Oxford, Oxford, OX3 7LE, UK

**Keywords:** Key Words: rheumatoid arthritis, circadian clock, inflammation, matrix metalloprotease

## Abstract

Rheumatoid arthritis is a chronic inflammatory disease that shows characteristic diurnal variation in symptom severity, where joint resident fibroblast-like synoviocytes (FLS) act as important mediators of arthritis pathology. We investigate the role of FLS circadian clock function in directing rhythmic joint inflammation in a murine model of inflammatory arthritis. We demonstrate FLS time of day-dependent gene expression is attenuated in arthritic joints, except for a subset of disease-modifying genes. Deletion of essential clock gene *Bmal1* in FLS reduced susceptibility to collagen-induced arthritis (CIA), but did not impact symptomatic severity in affected mice. Notably, FLS *Bmal1* deletion resulted in loss of diurnal expression of disease-modulating genes across the joint, and elevated production of MMP3, a prognostic marker of joint damage in inflammatory arthritis. This work identifies the FLS circadian clock as an influential driver of daily oscillations in joint inflammation, and a potential regulator of destructive pathology in chronic inflammatory arthritis.

## Introduction

Rheumatoid arthritis (RA) is a chronic, debilitating disease which affects 0.5-1% of adults in Western populations and requires long-term management^1^. Autoimmune activation leads to immune cell infiltration to the joint and inflammation of the synovial membrane. This results in local and systemic production of inflammatory cytokines and disease effector proteins including matrix metalloproteases, and subsequent progressive destruction of articular cartilage and bone. The associated pain, swelling, and loss of mobility have profound impacts on quality of life. Individuals with RA typically show time of day differences in the severity of these symptoms, associated with diurnal rhythms in inflammatory mediators and circulating cytokines^2^. The mechanisms and cell types which drive this strong daily rhythmicity in disease activity remain to be fully defined. The circadian biological clock has been strongly implicated^3–5^, and disruption of internal timing by lifestyle factors such as shift work is associated with elevated disease incidence^6^.

The circadian system orchestrates daily rhythmic fluctuations across much of mammalian behavior and physiology. At a molecular level, entraining signals (such as light, activity and nutrition) regulate rhythmic expression of core clock transcription factors (TFs). These TFs interact through coordinated transcription-translation feedback loops that cycle with a near 24h period^7^. The core circadian TFs additionally regulate numerous downstream target genes, giving rise to daily rhythms in cellular and physiological processes including metabolism, immune homeostasis and inflammatory response. This internal daily rhythm impacts disease processes involved in chronic inflammatory conditions, such as asthma and arthritis^8,9^.

Murine collagen-induced arthritis (CIA) is a well-established model of autoimmune, destructive, chronic arthritis, provoked by immunisation against type II collagen^10^. Our previous work has demonstrated the circadian rhythmicity of human disease is recapitulated in the CIA model, with clear time of day dependent fluctuation in the severity of joint inflammation^4,11^. Here, inflammatory gene expression in the joint and the abundance of associated circulating cytokines both peak during the rest phase for nocturnal mice, in agreement with human symptomatic peaks in the early morning. Our studies have also revealed that CIA-affected animals exhibit robust rhythmic gene expression in inflamed joint tissue, but this rhythmic transcriptional profile is profoundly different from that observed in naïve joints^11^. Interestingly, analyses of immune cell infiltration into the diseased joints reveals only limited time-dependent differences in cell number and subtype, as well as pronounced suppression of core clock function in infiltrating myeloid cells^12^. This suggests that rhythmicity within the inflamed joint is driven by other joint-resident cell populations.

Fibroblast-like synoviocytes (FLS) are mesenchymal-derived cells that reside in the joint synovial space and have important roles in the maintenance, repair and immune homeostasis of healthy joints^13,14^. During inflammatory arthritis, FLS undergo extensive functional remodelling, and drive tissue damage and inflammatory pathology. A number of FLS subpopulations have been identified in humans and mice^15–17^, with distinct cell surface marker expression profiles and phenotypic properties. In chronic joint disease, lining layer (LL) FLS are typically associated with expression of genes which contribute to destructive processes in the joint, whilst sub-lining layer (SLL) cells are associated with production of inflammatory signals and immune cell recruitment^15^. We have previously shown that FLS exhibit a robust internal circadian clock, and that modifying clock function can alter their inflammatory response^3,4^. These properties prompted us to hypothesise that joint-resident FLS may be a cellular driver of daily rhythms in inflammatory arthritis.

Here, we investigated FLS rhythmicity under normal conditions and in the context of CIA. Transcriptional profiling of FLS from naïve and CIA-affected joints revealed a pronounced attenuation of time of day-dependent gene expression in response to disease. Nevertheless a small, but potentially influential, subset of disease-related genes became robustly rhythmic in these cells. Upon deletion of the essential clock gene *Bmal1* in *Col6a1-*expressing FLS, we found a reduced incidence of CIA. However, in animals that progressed to severe disease, FLS-selective deletion of *Bmal1* led to a loss of time of day variation in markers of disease activity across the whole joint, supporting an important role for the FLS clock in the modulation of inflammatory processes and daily rhythms in joint disease.

## Results

### FLS exhibit altered rhythmic activity during CIA

To examine rhythmic transcriptional processes in FLS, we isolated Podoplanin (Pdpn)-positive cells from naïve and inflamed paws of wildtype DBA/1 mice at the peak and trough of disease activity (mid-light/rest phase, zeitgeber time ZT4, and mid-dark/active phase, ZT16; Fig. 1A-B). We found clear changes in gene expression in response to disease, with >8000 genes showing significant differential expression (DE) at each time point (Fig. 1C, Supplementary Dataset S1). Pathway analysis identified an enrichment of DE genes associated with KEGG pathways including ‘*rheumatoid arthritis*’ and ‘*cytokine-cytokine receptor interaction*’ (Fig. S1A, Supplementary Dataset S1), as would be expected. Time of day comparisons identified 1276 DE genes in FLS isolated from naïve mice, highlighting that these joint resident cells exhibit significant diurnal variation in function under homeostatic conditions. Gene enrichment analyses included pathways associated with ‘*steroid biosynthesis’* and ‘*circadian rhythm*’ (Fig. 1C, Supplementary Dataset S1). In contrast, we identified only 17 genes with significant DE between ZT4 and ZT16 in FLS isolated from inflamed joints (Supplementary Dataset S1). This profound loss of time of day dependent gene expression in response to CIA was not due to increased intragroup variability in samples isolated from CIA mice, or to a relatively minor reduction in the magnitude of expression change between time points in naïve and CIA derived FLS (Fig. 1D; Fig. S1B). This suggests a genuine and pronounced loss of rhythmic gene expression within FLS in the inflamed joint.

**Figure 1.**
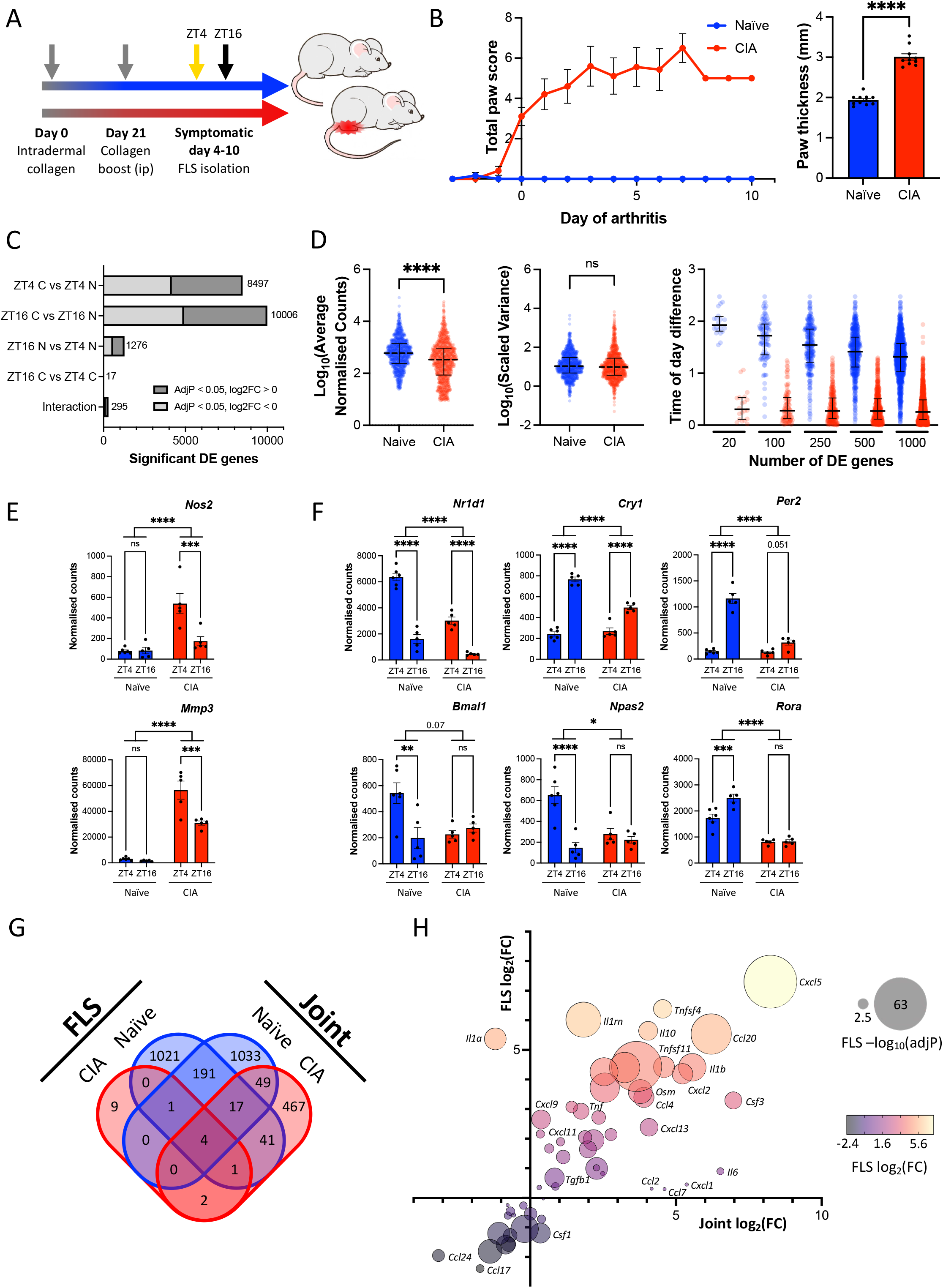
FLS exhibit altered rhythmic activity during CIA. **A.** Schematic representation of collagen-induced arthritis (CIA) experimental plan. **B.** Total paw score and paw thickness (day of sample collection) measurements of mice used for FLS isolation. Mean +/-SEM, Welch’s t-test, n = 10-11/condition. **C.** FLS cells from CIA or naïve mice were isolated at ZT4 or ZT16 and analysed by RNAseq (n = 5-6/time/condition). Differential expression analysis identified transcripts which are significantly up-or downregulated with disease and/or time. **D.** Peak expression level (left) and scaled variance (centre) were compared for the top 1000 significantly differentially expressed genes in naïve or CIA FLS cells, and the magnitude of time of day difference calculated (right; difference in mean expression level at ZT4 and ZT16 after z-scoring for top 1000 genes rhythmic in naïve FLS). Error bars show median +/- interquartile range, distributions compared by Welch’s t-test. **E.** *Nos2* and *Mmp3* show time of day dependent expression in FLS cells from inflamed joints. Mean +/- SEM, two-way ANOVA treatment effect and time-dependent differences are indicated. n = 5-6. **F.** FLS core clock gene expression in cells isolated from inflamed paws. Mean +/- SEM, two-way ANOVA, **G.** Four-way Venn comparison of time of day dependent gene expression between ZT4 and ZT16 timepoints in FLS RNAseq and whole joint RNAseq datasets^11^. **H.** Comparison of cytokine/chemokine expression changes with CIA in joints and isolated FLS cells. Of 92 transcripts considered (CCL, CXCL, interleukin, interferon, TNF and TGF families), 58 were detected in both sample types. Fold changes in expression were calculated from RNAseq data at ZT4.

The group of genes which showed significant time of day difference in FLS isolated from arthritic joints included a number encoding disease effector proteins including *Nos2* and *Mmp3* (Fig. 1E). These showed increased expression in CIA, with a notable emergence of time of day variation in expression, peaking at maximal disease activity (ZT4). These disease effectors have both previously been identified as being produced by synovial cells and chondrocytes in arthritis, with contributions to joint remodelling^18–21^. Whilst failing to reach our stringent FDR criterion, a number of other inflammation-associated chemokines and disease effector genes, including *Tnfsf11* (RANKL), *Mmp13, Ccl20* and *Cxcl5*, were significantly upregulated in CIA FLS on *post hoc* analysis, with higher expression at the peak of disease activity (Fig. S1C). Genes that remained robustly rhythmic in FLS from arthritic joints included circadian clock factors *Nr1d1* and *Cry1* (Fig. 1F). Interestingly, the impact of CIA was not uniform across the components of the molecular clock; genes associated with the negative regulatory arm of the clock (e.g. *Nr1d1*, *Cry1, Per2*) continued to show clear temporal expression, albeit at a reduced level, but positive regulators (e.g. *Bmal1*, *Npas2*, *Rora*) did not show time of day dependent expression in CIA (Fig. 1F).

We have previously reported that CIA results in a profound alteration to gene expression in whole joints, including emergence of a broad rhythmic transcriptional profile^11^. As this contrasts with our observed FLS gene expression patterns, we compared FLS-specific and whole joint transcriptional profiles in response to disease and time of day. We found that fewer than 10% of genes exhibiting time-dependent expression (significant DE in at least one condition) were common to FLS and whole joint data sets (Fig. 1G, Fig. S1D). Indeed, almost 600 genes showed time of day differences with CIA in bulk joint samples (Fig. 1G), including mediators of inflammatory signalling (*Stat3*) and cytokines (*Cxcl1, Il6, Ccl2*). We interpret this as representing time of day variation in expression of genes by non-FLS populations in the inflamed joint (e.g. immune cells). In support, comparison of FLS versus bulk joint expression of these rhythmic cytokines demonstrated limited disease-induced expression within FLS but substantial increase in expression within the joint as a whole (Fig. 1H). For example, *Il6* and *Cxcl1* are highly rhythmic and strongly induced in the whole joints (Fig. S1E), but this is not replicated in purified FLS, where CIA-induced expression is relatively limited. Of note, IL6 levels are also highly rhythmic in the circulation of both arthritic mice and humans^11,22^. This suggests that FLS are not the primary source of these signals.

### Loss of molecular clock function alters FLS transcriptional profile

Although FLS lost substantial transcriptional rhythmicity in arthritic joints, some important factors exhibited robust rhythmicity in these cells, including mediators which may be involved in communicating circadian timing and inflammatory state to other cell populations within the joint. We next assessed the role of FLS circadian oscillator function in disease pathogenesis using transgenic mice with selective deletion of the essential core circadian gene *Bmal1* in mesenchymal cells, including FLS, using the established *Col6a1*-cre driver line^3,23^. The resulting mice (*Col6a1^Cre+^Bmal1^fl/fl^*) were backcrossed onto a DBA/1 background to promote susceptibility to the CIA model of inflammatory arthritis. Targeting was confirmed in FLS isolated from *Cre* expressing mice (*Cre+*), which showed significant reduction in *Bmal1* transcript and protein expression compared with *Bmal1^flfl^* littermate controls (*Cre*-; Fig. 2A-B).

**Figure 2.**
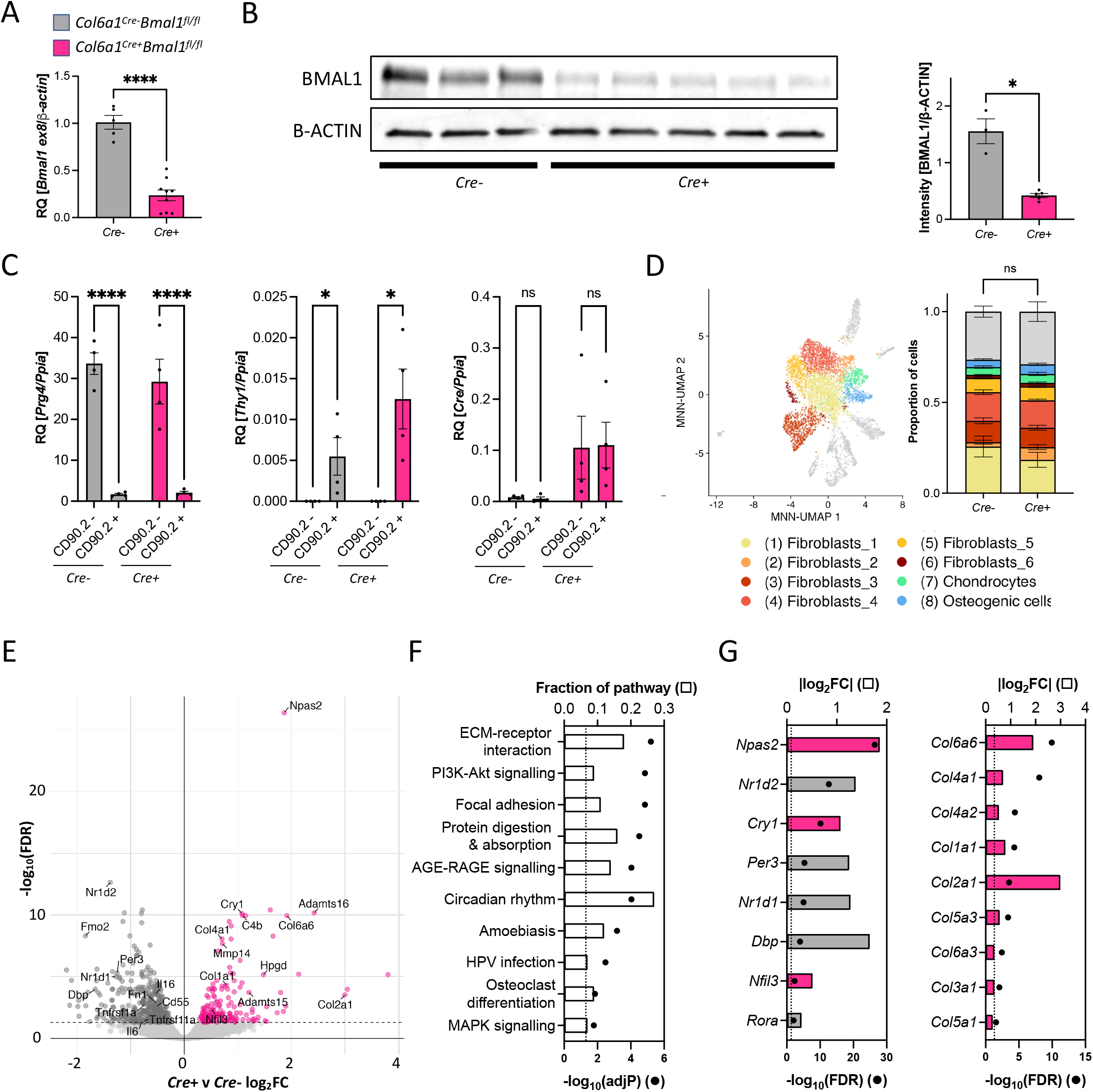
Loss of molecular clock function alters FLS transcriptional profile. **A, B.** Quantification of BMAL1 ablation at transcript (**A**) and protein (**B**) levels for *Cre*-and *Cre*+ mice. Mean +/- SEM, Welch’s t-test, n = 3-7/genotype. **C.** FLS cells sorted into CD90.2 negative and CD90.2 positive subpopulations were analysed to confirm lining layer (LL) versus sublining layer (SLL) identity (*Prg4* and *Thy1* marker gene expression, respectively; left, centre) and comparable expression of *Cre* recombinase (right). Gene expression is presented as mean +/- SEM, RM two-way ANOVA, post-hoc subpopulation comparison, n = 4 independent replicates. **D.** Gene expression of FLS isolated from joints was analysed using SPLIT-seq. Cells were clustered (left) and subpopulation identity determined based on cluster-specific expression of highly variable genes, identifying 6 FLS subpopulations (as well as chondrocytes, osteogenic cells and minor populations of other cell types), with no significant difference in population distribution between genotypes (right); two-way ANOVA, n = 3/genotype. **E.** Volcano plot showing differential gene expression between naïve *Cre*-(grey) and *Cre+* (pink) samples. Comparative expression analysis using edgeR considered pseudobulk populations of cells from clusters 1-8. **F.** Analysis of genes showing significant differential expression between genotypes identified enriched KEGG pathways (adjP < 0.05, dashed line). Fraction of pathway indicates the proportion of genes associated with a pathway which showed significant differential expression between genotypes. **G.** Circadian (left) and collagen/cell matrix-associated genes (right) are differentially expressed between genotypes (FDR < 0.05, dashed line). Differential expression is presented as the absolute value of log_2_ fold change, where grey indicates increased expression in *Cre-* mice and pink indicates increased expression in *Cre+* mice.

To ensure that *Col6a1*-cre targeting was effective across FLS subpopulations, we analysed gene expression in CD45^-^CD31^-^PDPN^+^ cells sorted based upon CD90.2 expression, a marker that allows distinction between sub-lining layer (SLL; CD90.2^+^) and lining layer (LL; CD90.2^-^) FLS. We found comparable expression of *Cre* recombinase in sorted LL and SLL cell populations from *Cre+* mice (Fig. 2C). In addition, we crossed *Cre+* mice with a line of mice that provide *Cre*-dependent EYFP expression, and found similarly robust EYFP fluorescence in LL and SLL cells (Fig. S2A). These data demonstrate effective targeting by the *Col6a1*-cre driver across FLS subpopulations.

We next used single cell RNA sequencing (scRNAseq) to characterise joint-derived PDPN^+^ cell identity and determine the impact of *Bmal1* deletion on FLS transcriptional phenotype. We identified multiple FLS subpopulations, and only minor contamination by other cell types (Fig. 2D, S2B-D). Overall, cellular subpopulation distributions in *Cre+* and *Cre*-FLS were broadly similar (Fig. 2D) and in line with previous work^14,16,17^, indicating that *Bmal1* expression is not required for differentiation of FLS subsets. We used a pseudobulk differential expression analysis approach to assess the impact of *Bmal1* deletion across all FLS populations. This identified 603 DE genes between *Cre+* and *Cre-* FLS (Fig. 2E, Supplementary Dataset S2). Ontology analysis identified a number of enriched KEGG pathways, including ‘*extracellular matrix (ECM)-receptor interactions’* and ‘*circadian rhythms’* (Fig. 2F, Supplementary Dataset S2). In line with this, FLS cells lacking *Bmal1* exhibited the expected altered expression of core clock genes (Fig. 2G). We also observed altered expression of numerous ECM factors including collagens (Fig. 2G), highlighting a potential role for the FLS clock in regulating ECM dynamics in the healthy joint.

### Mice lacking *Bmal1* expression in FLS show altered CIA development and loss of rhythmic joint inflammation

To understand the impact of clock ablation on disease phenotype, we next characterised the development and severity of CIA in *Cre+* mice and *Cre*-littermate controls. Interestingly, we found a significant reduction in the incidence of disease in *Cre+* mice (Fig. 3A, Fig. S3A-B). This was despite comparable anti-collagen IgG1 and IgG2a antibody responses, confirming the ability of both genotypes to mount a competent immune response to collagen immunisation (Fig. 3B). The cause of the reduced susceptibility to symptomatic disease is unclear, but may be due to cellular or structural changes within the joint. Our earlier work in C57BL/6J mice showed that similar deletion of *Bmal1* in *Col6a1*-expressing cells leads to age-related joint stiffening and increased chondrocyte number and paw thickness^3^. Here, we also observe a small but significant increase in paw thickness in naive *Cre+* mice compared to age-matched *Cre-* littermate controls (Fig. S3C). Elevated collagen gene expression by FLS which lack *Bmal1* expression (Fig. 2G) may contribute to both altered joint matrix and antigen response.

**Figure 3.**
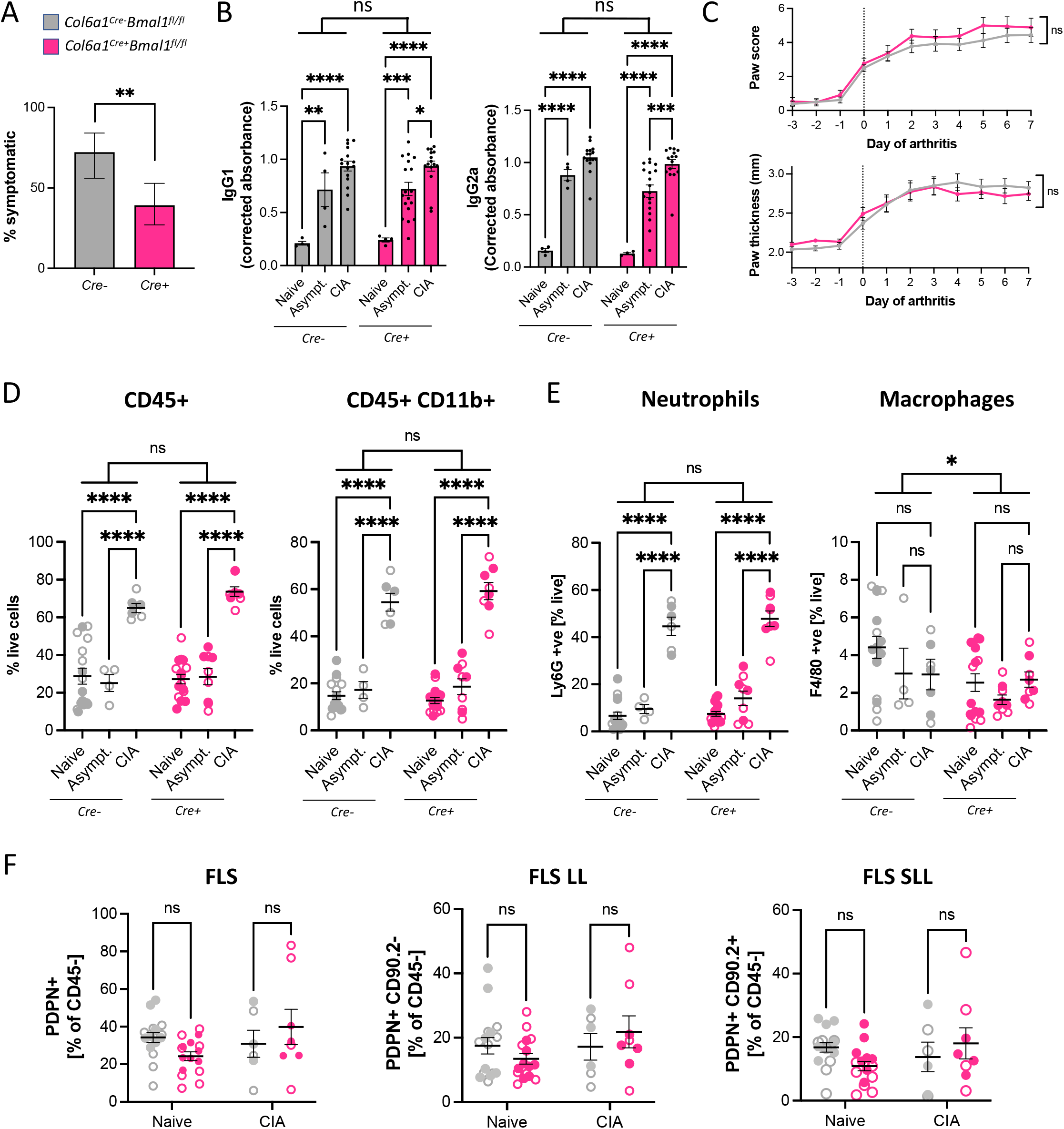
Mice lacking *Bmal1* expression in FLS show altered CIA development. **A.** Development of symptoms in *Cre*+ mice was significantly reduced compared to littermate *Cre*-controls. Fisher’s exact test; error bars indicate 95% confidence interval. **B.** Quantification of IgG1 and IgG2a anti-collagen antibody response in plasma from naïve, asymptomatic and symptomatic CIA mice. Two-way ANOVA, post-hoc treatment comparison, n = 4-18/condition. **C.** Daily paw score (top) and thickness measurement (bottom) found no difference in disease severity between mice that develop symptoms. Two-way ANOVA, mean +/- SEM, n = 17-26 mice/genotype. **D, E, F**. Flow cytometry analysis of immune cell infiltration into joints with disease, comparing total immune cell proportion (**D**) and cell identity (**E**) between genotypes. Flow cytometry analysis of FLS identity (**F**) comparing lining layer (LL) versus sublining layer (SLL) between disease and time of day. Open circles indicate ZT8 sample collection, filled circles indicate ZT20 sample collection. Two-way ANOVA, post-hoc treatment (**D**, **E**) or genotype (**F**) comparison, n = 3-8/time/condition.

Importantly, no significant differences were observed in markers of disease severity (paw score, paw thickness) between *Cre+* and *Cre-* animals which went on to develop arthritis (Fig. 3C). Therefore, to determine the impact of FLS *Bmal1* deletion on established joint disease, our subsequent analyses compare joints from symptomatic CIA mice with severe disease (paw score of ≥3). In comparison to joints isolated from naïve and asymptomatic mice, CIA caused a significant increase in infiltrating immune cells (predominantly neutrophils); this was similar between *Cre+* and *Cre-* genotypes, and did not show a time of day difference (Fig. 3D-E). Moreover, no genotype differences were observed in PDPN^+^ cell number (as a fraction of non-immune cells), the proportion of LL and SLL cells, or the expression of disease-associated FLS activation markers including CD106 (VCAM1) and FAP⍺ in naïve or CIA paws (Fig. 3F; Fig. S3D). Taken together, these data suggest that once disease is established, *Cre+* mice exhibit a similar profile of immune cell and FLS activation within affected joints to their control littermates.

To determine whether intrinsic clock function of FLS contributes to setting wider daily rhythms in inflammation across the whole joint, we isolated RNA from inflamed (CIA) and naïve whole joints of *Cre+* mice and *Cre-* littermate controls. We observed the expected significant elevation in proinflammatory gene expression in CIA joints; however, time of day variation in expression of key inflammatory cytokines (*Cxcl1, Il6, Il1b*) was found in *Cre-* but not *Cre+* joints (Fig. 4A). This suggests that the FLS clock is necessary to confer rhythmicity. Since we find limited expression of these cytokine transcripts in FLS themselves (Fig. 1H), this implicates FLS as orchestrators of disease rhythmicity across other resident or infiltrating cells.

**Figure 4.**
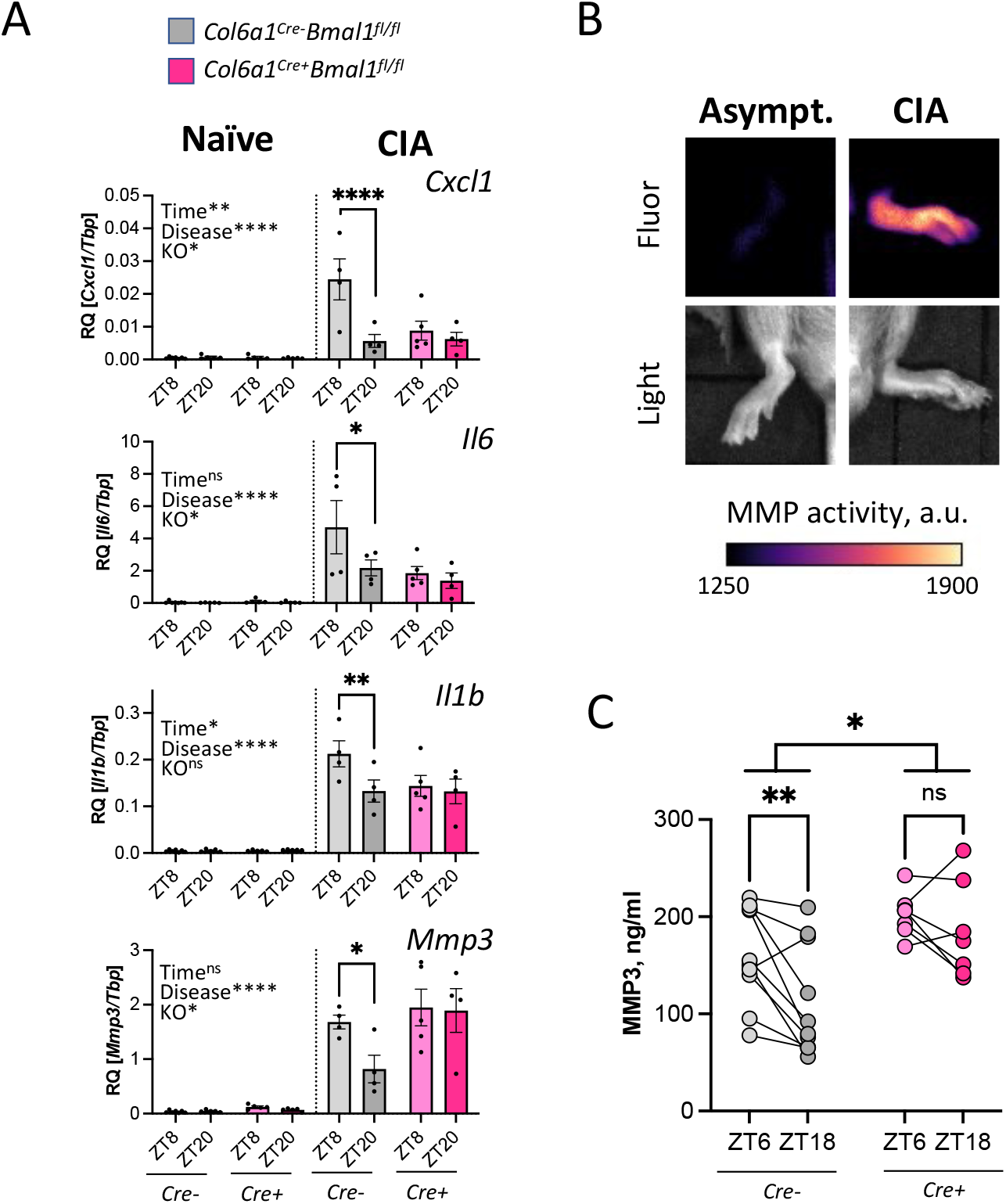
Mice lacking *Bmal1* expression in FLS lose rhythmicity of disease effectors in joints. **A.** Inflammatory gene expression in naïve and CIA joints from *Cre*-and *Cre*+ mice. Three-way ANOVA results and time-dependent differences in gene expression are indicated, n = 4-7/condition. **B.** Administration of an MMP activity-dependent fluorescent probe identified localised enzymatic activity in inflamed (but not asymptomatic) CIA paws. **C.** MMP3 level in plasma from arthritic mice across time. RM two-way ANOVA with post-hoc time comparison, n = 7-10/genotype.

CIA joints from *Cre-* mice exhibited a significant induction and time of day variation in *Mmp3* transcript (Fig. 4A). This time of day expression difference of *Mmp3* was lost in *Cre+* joints, with elevated expression at the typical nadir of inflammation. Significant MMP activity can be observed in symptomatic CIA paws *in vivo* (Fig. 4B, S4A). *Mmp3* expression was also persistently upregulated in FLS cultured from CIA paws (Fig. S4B) and strongly induced in cultured FLS by pro-inflammatory cytokine treatment (Fig. S4C). Analysis of plasma samples from CIA mice around the peak and nadir of arthritic inflammation revealed time of day differences in MMP3 levels at ZT6 compared to ZT18 in *Cre-* mice (Fig. 4C). In contrast, *Cre+* mice exhibited loss of time of day difference and augmented levels of MMP3 in plasma. Overall, this supports clock-dependent regulation of joint pathological processes by FLS.

## Discussion

RA, like many chronic inflammatory conditions, shows a characteristic variation in severity by time of day. Understanding the interplay between autoimmune inflammation and internal biological timing offers an opportunity to improve the management of chronic disease and identify new therapeutic approaches. Our previous work demonstrated damping of core clock gene rhythmicity in major immune cell populations harvested from arthritic joints, despite robust oscillations in joint transcriptional activity and circulating cytokine levels^11,12^; hence we hypothesized FLS, resident within the joint, might be a key source of these rhythmic inflammatory signals. FLS are a heterogenous population of cells that are intimately involved in the pathogenesis of RA^24–27^. Here, we find that diurnal rhythmicity in FLS gene expression is also suppressed by chronic inflammation. However, a subset of disease-relevant genes gain significant rhythmicity in these cells. Genetic disruption of *Bmal1* within FLS prevents time of day-dependent variation of key arthritis effector molecules within inflamed joints. Together, these studies identify an important functional role for the clock within FLS in promoting rhythmic signatures associated with chronic inflammatory arthritis.

A suppressive effect of inflammation and inflammatory cytokine signals on circadian gene expression has been widely reported *in vivo*^11,28–30^ and *in vitro*^31,32^. Transcriptional profiling of joint FLS from naïve and CIA mice showed a widespread damping of time of day dependent gene expression in these cells in response to arthritis. In contrast, prior whole joint sequencing revealed emergent inflammatory gene rhythmicity in diseased joints (in line with rhythmic symptoms). Importantly, *Col6a1^Cre+^Bmal1^fl/fl^* murine CIA studies revealed that key rhythmic inflammatory signatures in whole joint were dependent on *Bmal1* expression in FLS. Thus, while FLS are not the cellular source of the inflammatory rhythmic signature in arthritic joints, clock function within these cells appears to be fundamental in driving these oscillations. Target cell types responsible for producing the rhythmic pro-inflammatory profile likely include cells of a myeloid lineage. Indeed, we have shown that despite significant damping of their own intrinsic cellular clock in an arthritic joint, myeloid cell pro-inflammatory outputs appear to be shaped by the daily changes in the surrounding joint environment^12^. In the current study, FLS sequencing has revealed candidate signals through which FLS may impose rhythmicity across the joint. These potentially include NO production via rhythms in *Nos2* expression and rhythmic MMP3 production, both previously linked to arthritis-associated joint damage and regulation of inflammation^18,33,34^, as well as other inflammation-associated disease effectors (*Cxcl5, Ccl20, Mmp13, Rankl/Tnfsf11*). Further work is required to fully elucidate the mechanisms and cell types involved.

MMP proteins cause irreversible damage to cartilage and bone, resulting in ECM degradation in the inflamed joint^21,33,35–38^. MMP3 showed time of day differences in arthritic DBA/1 mice, with highest expression at the peak of inflammatory symptoms. MMP3 shows similar time of day variation in human inflammatory disease contexts^39^. We find that the FLS clock restrains MMP3 production, leading to consistently high plasma concentrations in the absence of *Bmal1*. This is in line with previous studies showing altered MMP3 expression in global *Bmal1* KO mice^40^ and *Bmal1*-dependent regulation of MMP production by chondrocytes^41^. In addition, modulation of the clock factor and BMAL1 target NR1D1 can repress production of MMP3 by FLS and reduce cartilage and bone loss in CIA^42^, and there is evidence of direct regulation of MMP3 by NR1D1 in some cell types^43,44^. MMP3 is of particular clinical interest as a biomarker for inflammatory arthritis since serum level reflects synovial fluid level^45,46^, and is a prognostic indicator of joint damage in human RA^47–49^. Targeting FLS-derived disease effectors has shown synergistic benefit in combination with standard RA treatments in preclinical trials^50–53^, and therapeutic modulation of FLS function (rhythmic activity, MMP expression) may present an opportunity for diagnostic and therapeutic benefit.

Genetic disruption of core clock function has typically been associated with increased severity of inflammatory response^54–57^. Global deletion of *Nfil3*, *Cry1* and/or *Cry2* worsens disease severity in experimental arthritis models^58–60^. We have previously shown that in collagen antibody-induced arthritis (CAIA, a model which does not generate B and T cell-mediated autoimmune responses^61^), *Bmal1* deletion in FLS resulted in increased neutrophil infiltration and elevated inflammatory gene expression in affected joints^3^. In the CIA model of autoimmune mediated arthritis in the susceptible DBA/1 genetic background, *Bmal1* deletion in FLS led to a reduction in the incidence of disease compared with control littermates. This may reflect differences in genetic background of the mice, but more likely reflects differences in the immune mechanisms involved in disease induction between the two experimental arthritis models (CAIA and CIA). This would suggest that FLS, and *Bmal1* activity within these cells, are important in early immune priming events and/or initiation of the immune response at the joint synovium.

In a naïve state, *Col6a1^Cre+^Bmal1^fl/fl^* DBA/1 mice exhibit a small increase in paw thickness and enhanced FLS expression of collagens and other ECM-associated genes. This is in line with our previous work showing that ablation of *Bmal1* using the same *Col6a1*-cre driver in C57BL/6J mice results in joint stiffening^3^. *Bmal1* ablation in other tissue contexts has also been shown to alter ECM organisation^62,63^. Together these findings reveal a role for FLS BMAL1 activity or wider circadian clock function in shaping joint ECM dynamics, and we speculate that altered collagen expression and/or resulting joint architecture contribute to the attenuated susceptibility to CIA observed here. Indeed, several studies have found prior dosing with soluble collagen can reduce CIA incidence, whilst administration after the development of symptoms can increase anti-inflammatory cellular response and reduce expression of proinflammatory markers^64–66^.

In summary, we show that the FLS clock machinery is robust in healthy joints and directs a programme of rhythmic gene expression. However, that rhythmic activity is significantly attenuated during inflammation. Despite this attenuation, *Bmal1* expression in FLS is required for rhythmic expression of cytokines and other disease effector molecules in the arthritic joint. As we find FLS are not the cellular source of many of the cytokines typically associated with rhythmic symptoms in RA, this demonstrates the role of FLS in driving rhythmic function in other immune populations of the diseased joint.

## Acknowledgements

The work was supported by grants from the Medical Research Council (MRC; MR/P023576/1 to DWR, DAB, JEG; MR/V034049/1 to DWR, DAB; MR/W019000/1 to DWR), the Biotechnology and Biological Sciences Research Council (BBSRC; BB/V002651/1 to DAB), Versus Arthritis (22625 to JEG), the Wellcome Trust (107849/Z/15/Z to DWR), the NIHR Oxford Biomedical Research Centre (NIHR203316 to DWR) and a University of Manchester Facilitating Excellence Fund award to PD. We acknowledge the excellent experimental support of the University of Manchester Core Facilities: Dr Andy Hayes (Genomic Technologies), Dr Leo Zeef and Dr I-Hsuan Lin (Bioinformatics), and Dr Gareth Howell and Mr David Chapman (Flow Cytometry). We also thank and acknowledge the University of Manchester Biological Services Facility staff for animal care, and Dr Edward Hayter for assistance with *in vivo* experiments.

## Author Contributions

Conceptualisation, PD, DAB, JEG; Methodology, PD; Formal Analysis, PD; Investigation, PD, SHD; Writing – Original Draft, PD; Writing – Review & Editing, PD, DAB, JEG; Supervision, DAB, JEG; Project Administration, DWR, DAB, JEG; Funding Acquisition, DWR, DAB, JEG.

## Declaration of Interests

The authors declare no competing interests.

## STAR★Methods

### Key resources table

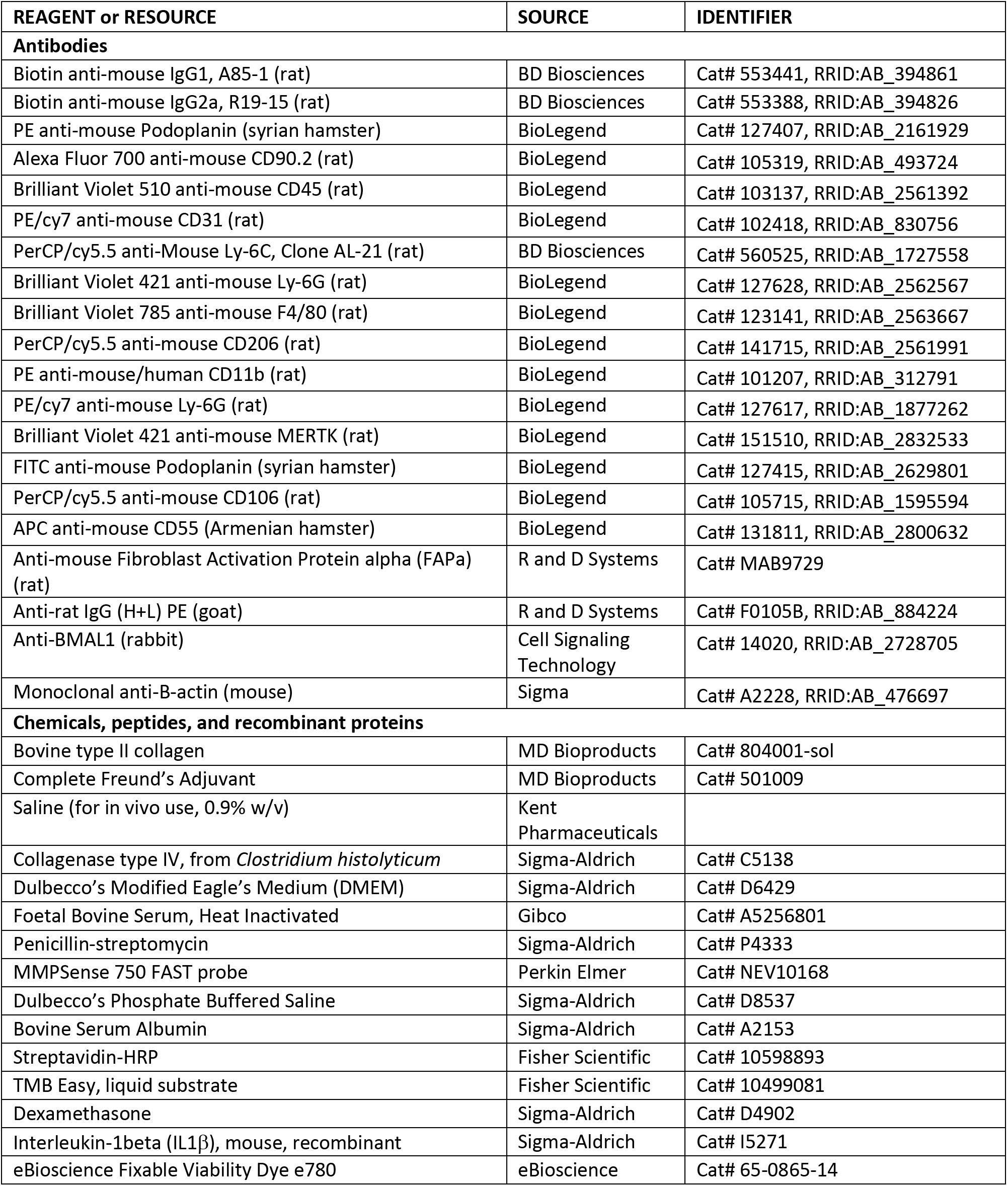
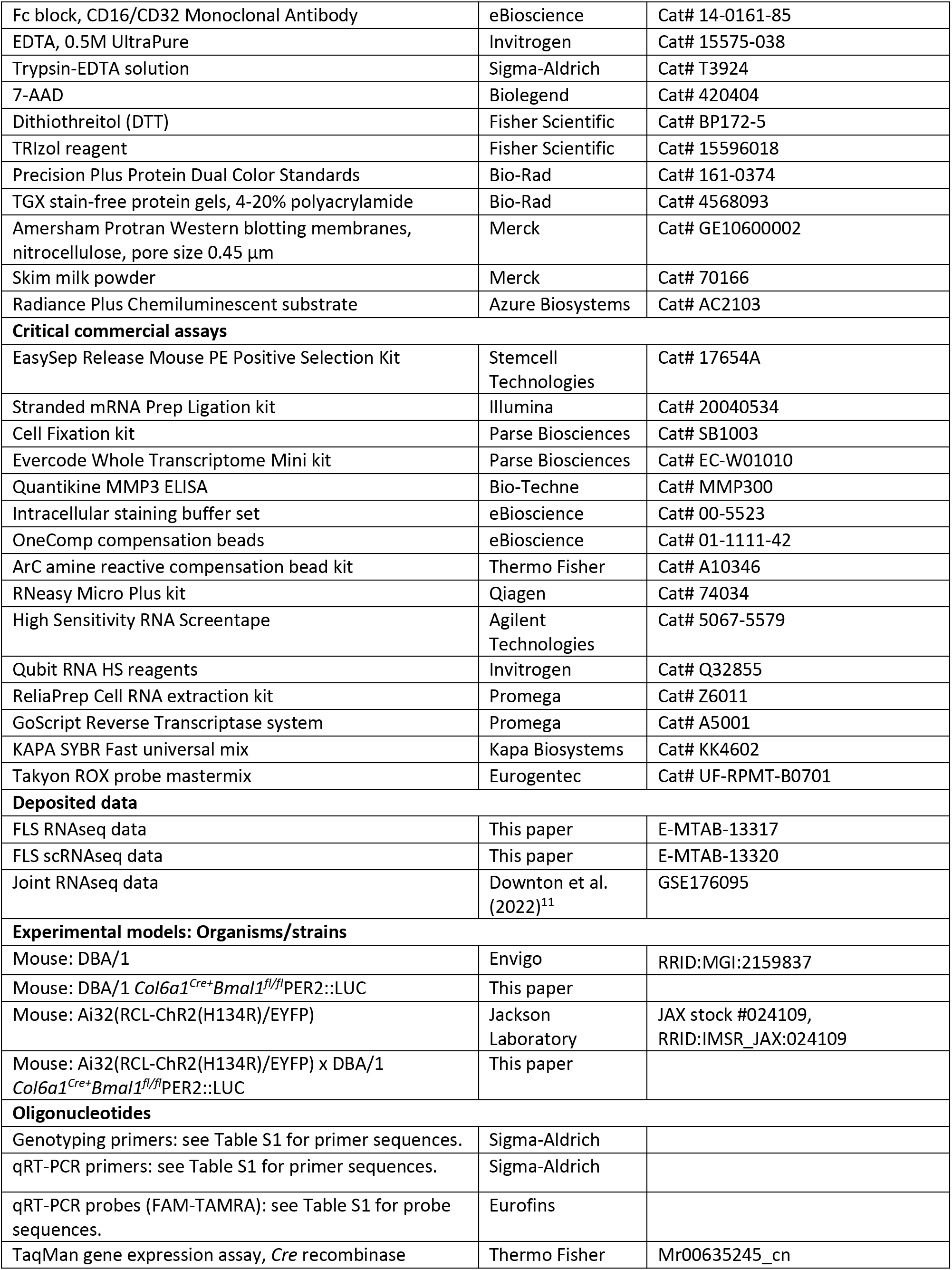
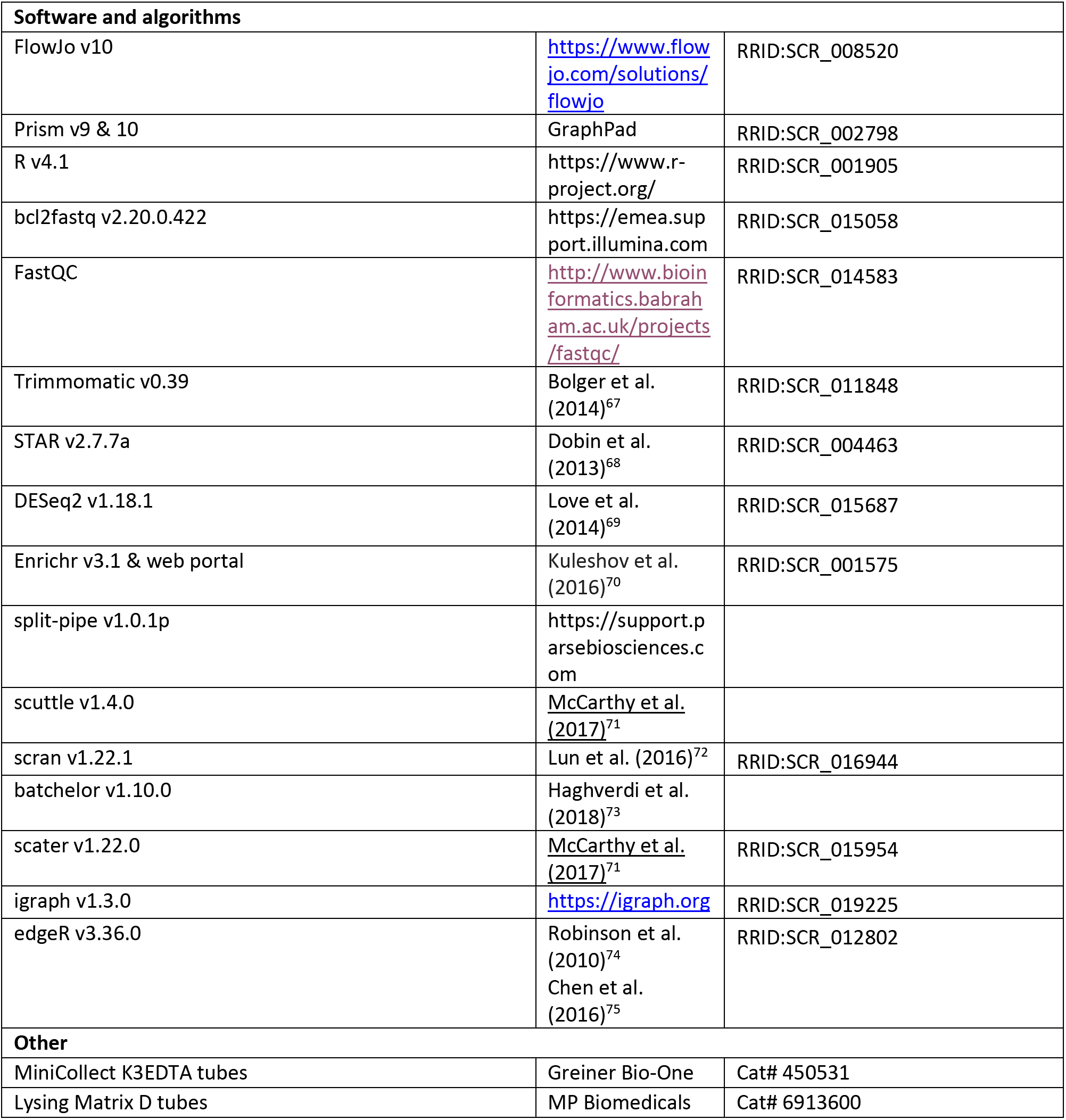

### Resource availability

#### Lead contact

Further information and requests for resources and reagents should be directed to and will be fulfilled by the lead contact, Professor Julie Gibbs.

#### Materials availability

Further information and requests for resources and reagents should be directed to and will be fulfilled by the lead contact, Professor Julie Gibbs.

#### Data and code availability

FLS RNAseq data from this study have been deposited in ArrayExpress with accession code E-MTAB-13317. scRNAseq data from this study have been deposited with accession code E-MTAB-13320. Published joint RNAseq data used for comparative analysis is available in Gene Expression Omnibus with accession code GSE176095.

Code will be available from GitHub upon publication.

Any additional information required to reanalyse the data reported in this paper is available from the lead contact upon request.

### Experimental model and study participant details

#### Animals

Mice were maintained in the University of Manchester Biological Services Facility. All procedures were approved by the University of Manchester Animal Welfare and Ethical Review Body and carried out according to the Animals (Scientific Procedures) Act 1986 under Home Office project licence numbers P000BBBC3 and PP5819195. Mice were group housed under a 12h:12h light/dark cycle with *ad libitum* access to food and water. DBA/1 mice were purchased from Envigo (Huntingdon, UK). Transgenic mice created for this study were generated in house. Male *Col6a1^Cre+^Bmal1^fl/fl^*PER2::LUC mice^3^ were backcrossed with female DBA/1 mice for three generations, then interbred to obtain homozygosity of the Bmal1^flox/flox^ modification. Ai32(RCL-ChR2(H134R)/EYFP) mice, which express a channel rhodopsin-EYFP fusion protein in Cre positive cells^76^, were crossed with DBA/1 *Col6a1^Cre+^Bmal1^fl/fl^*PER2::LUC mice for Cre-expression mapping experiments. Primers used for genotyping are given in Table S1.

#### Primary cell isolation and culture

Paws were dissected to separate joints, then incubated with collagenase IV (10 mg/ml, Sigma C5138) in DMEM containing 10% FBS and 1% penicillin-streptomycin at 37°C with shaking for 2 hours. Cells were maintained in DMEM containing 10% FBS and 1% penicillin-streptomycin in a humidified incubator (37°C, 5% CO_2_). Cells between passages 3-7 were used for all experiments. For *in vitro* experiments, cells were treated with recombinant mouse IL1β (final concentration 10 ng/ml, Sigma-Aldrich I5271-5UG).

### Method details

#### Collagen induced arthritis (CIA)

CIA was induced in male mice (7-13 weeks) as described previously^11^. Initial intradermal administration of bovine type II collagen in Complete Freund’s Adjuvant (MD Bioproducts, Zurich, 804001-sol and 501009) was followed by i.p. booster of collagen in saline on day 21. Disease severity and paw thickness were assessed daily. Disease severity was scored on a four-point scale for each paw (1 for single inflamed digit, 2 for multiple inflamed digits, 3 for swelling of paw pad, 4 for severe swelling of paw pad and ankle/wrist joint)^4^. Terminal tissue samples were collected 7-10 days after the development of symptoms unless otherwise specified.

#### *In vivo* procedures

Small volume blood samples were collected from the tail vein 5-7 days after the development of symptoms. Blood was collected into EDTA-coated tubes and plasma isolated by centrifugation at 3000 x *g* for 10 minutes at 4°C. For in vivo imaging studies, MMP activity was assayed using MMPSense 750 FAST probe (Perkin Elmer, NEV10168). Images were collected using an IVIS Lumina system (745 ex/800 em). After collection of a baseline image, the probe was administered i.v. (1.5 - 2 nmol), followed by image collection under isofluorane anaesthesia 12h after probe administration.

Terminal blood was collected in MiniCollect K3EDTA tubes (Greiner Bio-One) and stored on ice prior to centrifugation at 3000 x *g* for 10 minutes at 4°C. For whole joint RNA analysis, paws were dissected and skin removed prior to being flash frozen in liquid nitrogen. For FLS RNAseq and scRNAseq analysis, skin was removed and the paw incubated in cell culture media prior to joint digestion and FLS isolation.

#### FLS enrichment

Paws were dissected and incubated with collagenase IV (10 mg/ml, Sigma C5138) as described above. FLS for RNAseq and scRNAseq analysis were selected following 1-1.5h collagenase digestion using the EasySep Release Mouse PE Positive Selection kit (Stemcell Technologies, 17656) and a Podoplanin (PDPN)-PE conjugated antibody (BioLegend 127407). Efficiency of selection was tested by staining of released cells (or parallel unprocessed samples) for flow cytometry, and enrichment of the target cell population is shown in Figure S5A.

#### Flow cytometry and cell sorting

Joint cells for flow cytometry were isolated as described above, with 1-1.5h collagenase digestion. Cells were passed through a 40 µm filter, washed with PBS, then subjected to subsequent incubations with live/dead stain (20 minutes at 4°C, eBioscience Fixable Viability Dye e780, Invitrogen 65-0865-14), Fc block (20 minutes at 4°C, eBioscience 14-0161-85), and extracellular antibody mix (1h at 4°C, Table S2). Following extracellular antibody incubation, cells were washed once with FACS buffer (PBS containing 4% FBS and 10 mM EDTA) then fixed using an intracellular staining buffer set (eBioscience 00-5523). Cells were kept at 4°C until analysis, or subjected to intracellular antibody staining following overnight incubation in fixative. Where intracellular staining was necessary, cells were washed with permeabilization buffer, then incubated with primary intracellular antibody in permeabilization buffer (30 minutes at 4°C, Table S2). This process was repeated with a fluorescent secondary antibody if the primary antibody was unconjugated, and cells were resuspended in FACS buffer prior to analysis. In vitro cultured FLS cells were washed with PBS, detached using trypsin and resuspended in FACS buffer prior to staining using the same protocol. Stained cells were passed through a 40 µm filter and analysed on a BD LSRFortessa instrument. Compensation for spectral overlap used OneComp (eBioscience 01-1111-42) and ArC amine reactive (Thermo Fisher A10346) compensation beads. Data was analysed using FlowJo v10. Gating strategies for flow and cell sorting experiments were set using fluorescence minus one (FMO) control samples and are described in Figure S5B-D.

Joint cells for sorting were isolated as above, except the live/dead staining step was omitted. The antibody panel used for staining is described in Table S2. Cells were sorted into cold PBS using a BD FACSAria instrument. Immediately prior to sorting, 7-AAD (50 ng/ml, Biolegend 420404) was added to the filtered cell suspension. After sorting, cell pellets were collected by centrifugation and resuspended in buffer RLT Plus (Qiagen) prior to RNA isolation and analysis.

#### RNA isolation

RNA was extracted from PDPN-selected FLS cells and sorted FLS cells using the RNeasy Micro Plus kit (Qiagen). The FLS cell pellets were lysed in buffer RLT Plus supplemented with DTT, and RNA isolated according to the manufacturer’s instructions. RNA integrity was assessed using a 4200 TapeStation (Agilent Technologies).

RNA was extracted from cultured FLS cells using the ReliaPrep Cell kit (Promega). Cells were cultured in 24 or 12 well plates. After the indicated treatment, cells were washed in cold PBS then lysed by direct addition of buffer BL + TG to culture plate wells, followed by scraping and collection into microcentrifuge tubes. RNA was isolated according to the manufacturer’s instructions.

RNA was extracted from whole joint samples using TRIzol. Tissue was ground with liquid nitrogen then homogenised in a Lysing Matrix D tube using a BeadMill homogeniser (3 x 4 m/s for 40s). RNA was extracted using chloroform then precipitated with isopropanol. Isolated RNA was further purified using the RNeasy Micro Plus kit (Qiagen).

#### RNAseq

Sequencing library preparation and sequencing was performed by the University of Manchester Genomic Technologies Core Facility. Libraries were generated using the Stranded mRNA Prep Ligation kit (Illumina, Inc) according to the manufacturer’s protocol. Libraries were pooled prior to loading onto an S1 flow cell for paired end sequencing (59 + 59 cycles, plus indices) on an Illumina NovaSeq6000 instrument. The binary base call (BCL) files were processed using bcl2fastq software (v. 2.20.0.422).

#### scRNAseq

FLS cells were isolated using the EasySep Release PE positive selection kit and PDPN-PE antibody, as described above. Selected cells were fixed immediately after bead release using Cell Fixation reagents (Parse Biosciences SB1003) in accordance with the manufacturer’s instructions. Fixed cell suspensions were counted using a haemocytometer before being flash frozen in liquid nitrogen and stored at -80°C prior to barcoding and library preparation. Libraries for sequencing were prepared using the Evercode Whole Transcriptome Mini kit (Parse Biosciences EC-W01010). Briefly, fixed cells were barcoded by in cell reverse transcription, with samples distributed between 12 wells containing specific barcoded primers, followed by two subsequent rounds of mixing and ligation (2 x 96 wells containing specific barcodes) to generate unique combinatorial barcodes for each cell. Cells are then lysed and barcoded cDNA isolated, amplified and cleaned for sequencing library generation. Library fragment size and quality was assayed using a 4200 TapeStation (Agilent Technologies). Libraries were loaded onto an SP flow cell for paired end sequencing (74:86, c/w 5% PhiX spike-in) on the Illumina NovaSeq6000 platform, and BCL files processed as above.

#### Bioinformatic analysis of sequencing data

Bioinformatic analysis was undertaken in collaboration with the University of Manchester Bioinformatics Core Facility.

Sequencing quality was assessed using FastQC. FLS RNAseq data sequence adapters were removed, and reads were quality trimmed using Trimmomatic_0.39^67^. The reads were mapped against the reference mouse genome (mm10/GRCm38) and counts per gene were calculated using annotation from GENCODE M25 using STAR_2.7.7a^68^. Normalisation, Principal Components Analysis, and differential expression was calculated with DESeq2_1.18.1^69^. Adjusted p-values were corrected for multiple testing using the Benjamini and Hochberg method. Pathway analysis used the enrichR web portal to query the KEGG 2019 Mouse database^77^. For time of day/disease comparison of FLS and joint RNAseq datasets, previously published joint RNAseq data from samples collected at ZT4 and ZT16 were reanalysed using the same pipeline^11^.

FLS single cell FASTQ files were mapped against the Mouse reference GRCm39 and Ensembl annotation (v107) using the Parse Biosciences pipeline “split-pipe” (v1.0.1p) to generate the gene-cell count matrix. These were processed in R environment (v4.1) following the workflow of Amezquita *et al*^78^. Briefly, the “all-well” count matrix was imported into R to create a SingleCellExperiment object. A combination of median absolute deviation (MAD), as implemented by the “isOutlier” function in scuttle (v1.4.0) and exact thresholds were used to identify and subsequently remove low quality cells. The expression values were log-normalised and the per-gene variance of the log-expression profile was modelled using the “modelGeneVar” function and top 1500 highly variable genes (HVGs) were identified using the “getTopHVGs” function, both from scran (v1.22.1). Batch effects were corrected for visualisation using the mutual nearest neighbors (MNN) approach implemented by the “fastMNN” function from batchelor (v1.10.0). The MNN corrected coordinates were used as input to produce the uniform manifold approximation and projection (UMAP) using the “runUMAP” function from scater (v1.22.0) respectively. Cell subpopulations were clustered using the Leiden algorithm from igraph (v1.3.0). Marker genes were identified using the “findMarkers” function from the scran R package. Differential expression (DE) analysis between WT and KO populations was performed on pseudo-bulk samples from clusters 1 to 8 using the quasi-likelihood pipeline from edgeR (v3.36.0). Genes with a false discovery rate (FDR) below 5% were considered differentially expressed.

#### Quantitative PCR (qPCR)

RNA was converted to cDNA using the GoScript Reverse Transcriptase System (Promega A5001). Quantitative PCR used KAPA SYBR fast universal mix (Kapa Biosystems KK4602) or Takyon ROX probe mastermix (Eurogentec UF-RPMT-B0701) and a StepOnePlus Real-Time PCR machine (Applied Biosystems) with StepOne software v2.3. Primer sequences, probe sequences and Taqman assays are listed in Table S1.

#### Collagen antibody ELISA

Tissue-culture treated 96-well plates were coated by incubation with collagen (5 µg/ml, MD Bioproducts 804001-sol) at 4°C overnight. The plate was washed with PBS containing 0.1% Tween20 (PBS-T) then blocked with 2% BSA in PBS for 1 hour. Plates were washed with PBS-T, then serial dilutions of test plasma samples were added and left at room temperature for 1 hour. Plates were washed, incubated with a biotin-conjugated secondary antibody (BD Biosciences, A85-1 (anti-IgG1) and R19-15 (anti-IgG2a)) for 45 minutes, washed, incubated with streptavidin-HRP (Fisher Scientific 10598893) for 45 minutes, washed, then incubated with TMB Easy (Fisher Scientific 10499081). Reactions were stopped by addition of sulphuric acid, and absorbance read at 405 nm using a GloMax multi detection system (Promega).

#### MMP3 ELISA

MMP3 was quantified in serial plasma samples using a Quantikine ELISA kit (Bio Techne MMP300) according to the manufacturer’s instructions, with plasma diluted 1 in 25 in calibrant diluent. Plate absorbance was measured on a GloMax multidetection system (Promega).

#### Western blotting

FLS cells were lysed in preheated buffer (2% (w/v) SDS, 10% (v/v) glycerol, 50 mM DTT, 40 mM Tris pH 6.8, 0.001% (w/v) bromophenol blue). Protein samples and standards (Bio-Rad 161-0374) were resolved on TGX stain-free gels (Bio-Rad) run under denaturing conditions, then transferred to nitrocellulose membrane (Sigma-Aldrich GE10600002). Membranes were blocked with 5% (w/v) skim milk powder in TBS-T then incubated overnight with primary antibody (anti-BMAL1, 1/1000, Cell Signaling Technology 14020; anti φ3-ACTIN, 1/5000, Sigma-Aldrich A2228). Membranes were washed with TBS-T, incubated with HRP-conjugated secondary antibody, washed and developed using Radiance Plus chemiluminescent substrate (Azure Biosystems AC2103). Blots were imaged on a ChemiDoc imaging system (Bio-Rad). Uncropped Western blots are shown in Figure S6.

### Quantification and statistical analysis

Statistical tests were conducted in GraphPad Prism and are specified in figure legends where appropriate. Throughout, * denotes p<0.05, ** denotes p<0.01, *** denotes p<0.001 and **** denotes p<0.0001.

## Notes

### Competing Interest Statement

The authors have declared no competing interest.

