## Supplementary Information for "Fibroblast-like synoviocytes orchestrate daily rhythmic inflammation in arthritis"

**This PDF file includes:**

Figures S1 to S6
Tables S1 to S2
Legends for Supplementary Datasets

**Other supplementary materials for this manuscript include the following:**

Dataset S1
Dataset S2

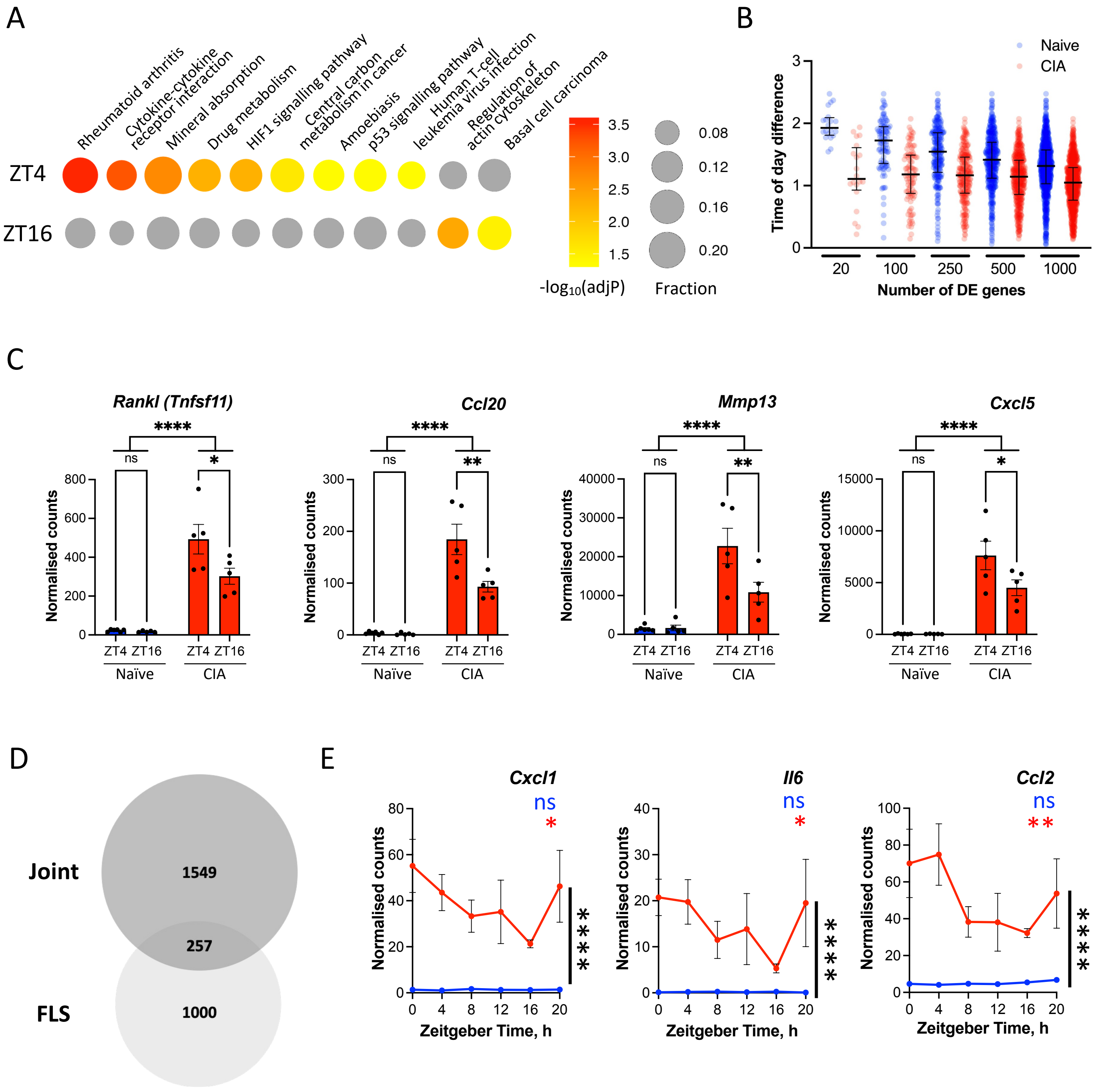


**Supplementary Figure S1.**

**Supplementary Figure S1**

**A**. Pathway enrichment analysis of genes differentially expressed (DE) with CIA in FLS at ZT4 and ZT16. KEGG pathways are shown if identified as significantly enriched at either ZT4 or ZT16; full analysis is provided in Dataset S1. Circle size indicates DE proportion of KEGG pathway; colour scale indicates significance of enrichment, grey indicates adjP > 0.05.

**B.** Z-scored time of day expression difference for the 1000 most significant DE genes in naïve (blue) and CIA (red) FLS.

**C.** Expression profiles for selected genes upregulated in CIA FLS. Mean +/- SEM, two-way ANOVA, post-hoc time comparison, n = 5-6/time/condition.

**D.** Comparison of DE genes that show time of day differences between ZT16 v ZT4 in whole joint or isolated FLS cells. Comparisons were made using the DEseq2 R package.

**E.** Whole joint expression profiles for selected genes upregulated in CIA. Mean +/- SEM, CIA vs naïve comparison by two-way ANOVA is indicated in black, adjP value of JTK rhythmicity analysis is indicated in blue (naïve) and red (CIA), n = 5/time/condition. *P < 0.05, **P < 0.01, ***P < 0.001, ****P < 0.0001.

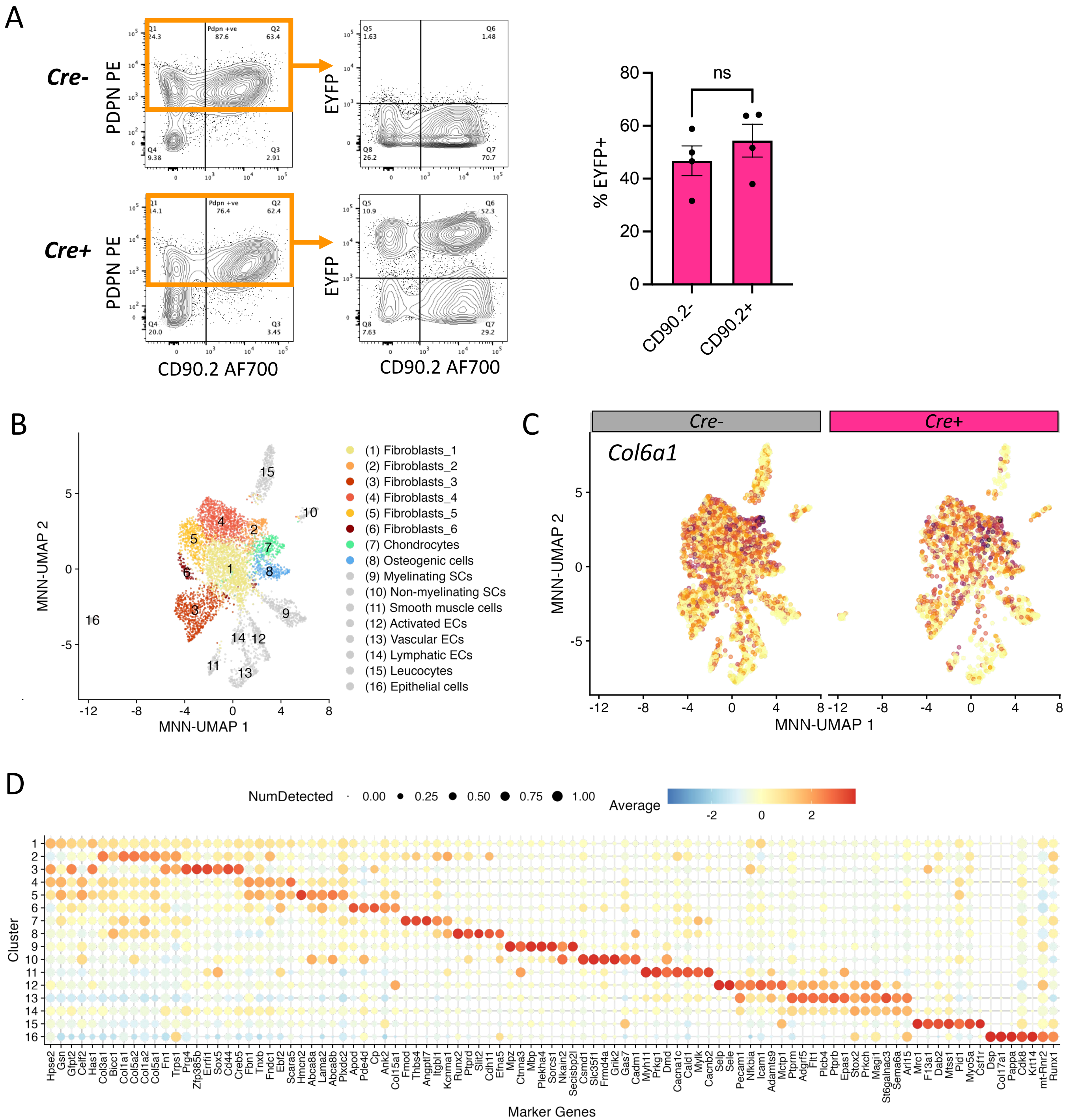


**Supplementary Figure S2.**

**Supplementary Figure S2**

**A.** Representative flow cytometry plots (left) showing EYFP expression in CD90.2 negative and positive FLS populations of *Cre*+ EYFP mice. Proportion of EYFP+ cells was not significantly different between CD90.2 negative and positive populations (right); paired t-test, n = 4/subpopulation.

**B.** UMAP plot representing subpopulations identified in single cell RNAseq experiment. Cell type was assigned based upon subpopulation expression of marker genes.

**C.** UMAP plots representing *Col6a1* expression in *Cre*- (left) and *Cre*+ (right) FLS.

**D.** Dot plot showing expression of cluster-specific differentially expressed genes for each identified subpopulation in **(C)**.

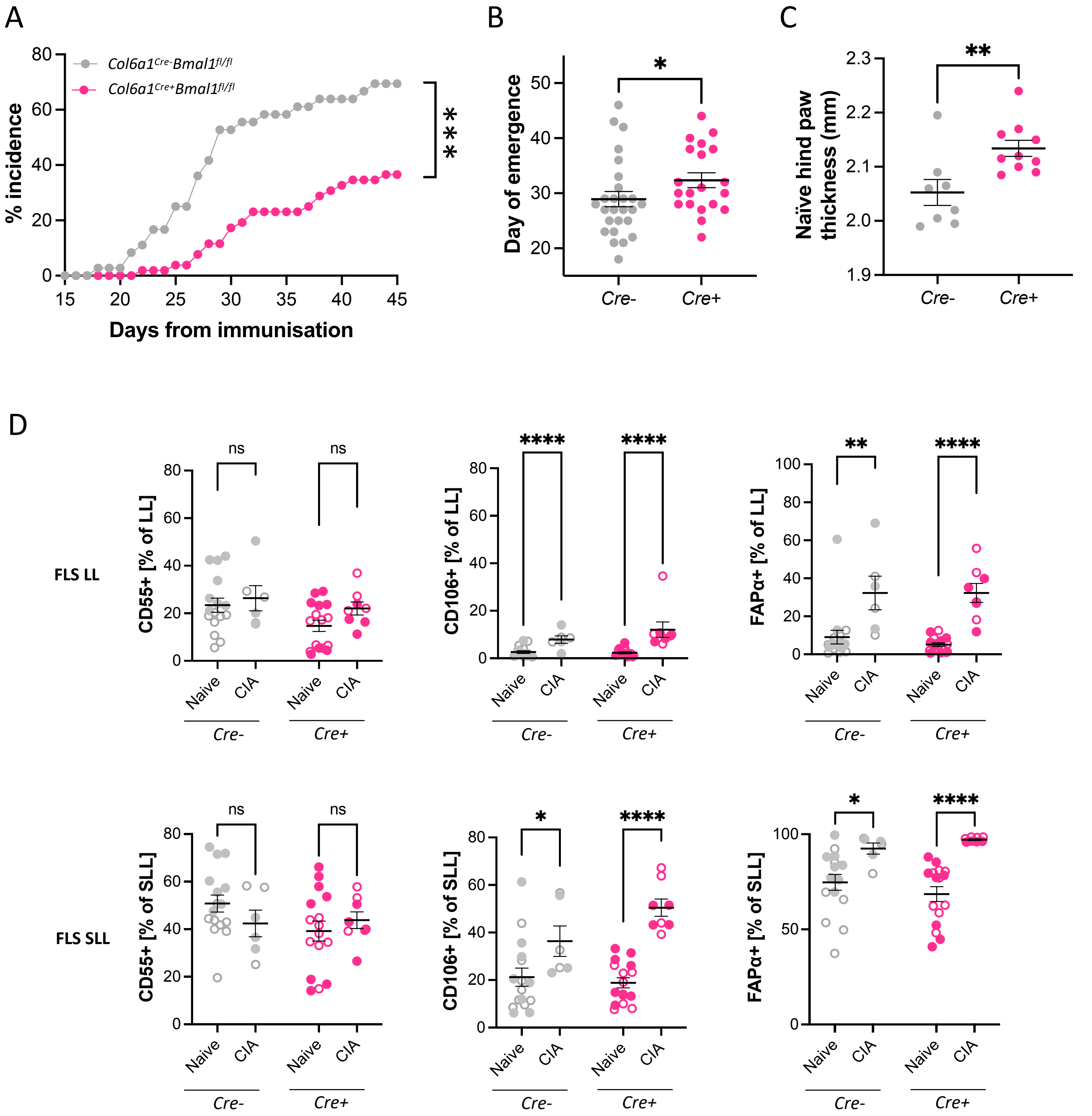


**Supplementary Figure S3.**

**Supplementary Figure S3**

**A.** *Cre*- (grey) and *Cre*+ (pink) disease emergence over the experimental period. Mice were considered to be symptomatic upon development of any clear disease symptoms within 45 days from initial collagen immunisation (paw score >0). Incidence was compared by log rank (Mantel-Cox) test, 1df, n = 36 *Cre*-, 52 *Cre*+ mice.

**B.** Day of symptomatic emergence in *Cre*- and *Cre*+ mice. Two-tailed Mann-Whitney test, n = 20-26 symptomatic mice/genotype.

**C.** Average naïve hind paw thickness on day of tissue collection. Mann-Whitney test, n = 8-10 mice/genotype.

**D.** Flow cytometry analysis of disease marker expression in LL and SLL FLS cells. Open circles indicate ZT8 sample collection, filled circles indicate ZT20 sample collection. Two-way ANOVA with post-hoc genotype comparison, n = 3-8/time/condition.

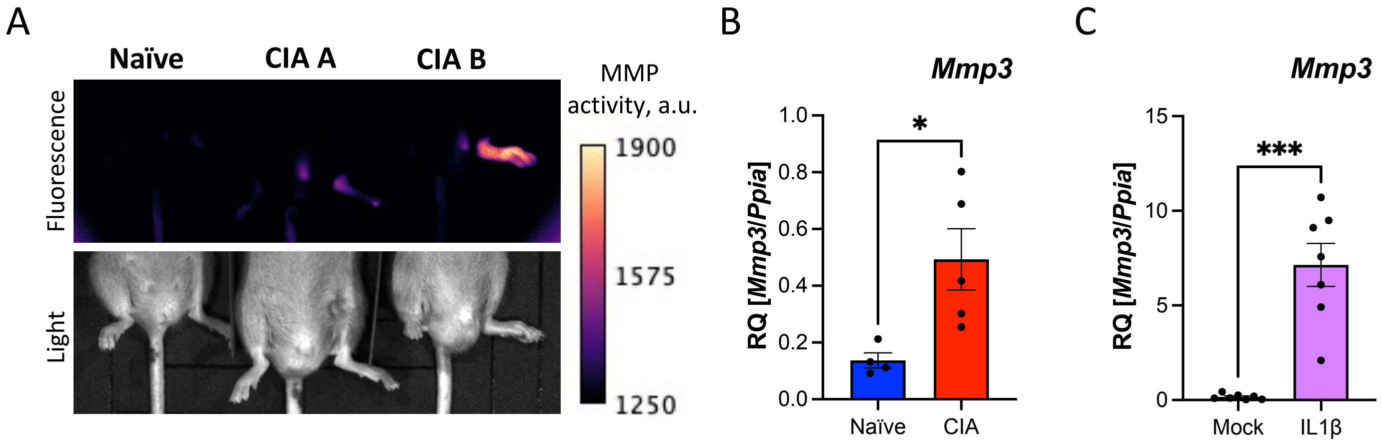


**Supplementary Figure S4.**

**Supplementary Figure S4**

**A.** In vivo fluorescent images acquired from mice administered an MMP activity-sensitive fluorescent probe. Probe was given 12 hours prior to image acquisition from one naïve (left) and two CIA (right) mice. Mouse CIA A exhibited paw scores of 1 (left paw; single digit inflamed) and 0 (right paw; asymptomatic) when imaged. Mouse CIA B exhibited paw scores of 4 (left paw; inflammation of the paw and ankle) and 0 (right paw; asymptomatic) when imaged.

**B.** Comparison of expression level of *Mmp3* in FLS isolated for culture from naïve or CIA joints. FLS were passaged 6 days after isolation and maintained in culture for 13-14 days prior to RNA isolation. Two-sided Mann-Whitney test, n = 4-5/condition.

**C.** Comparison of expression level of *Mmp3* in cultured FLS isolated from naïve mice. RNA was isolated following 48h cytokine stimulation or mock treatment. Paired t test, n = 7/treatment.

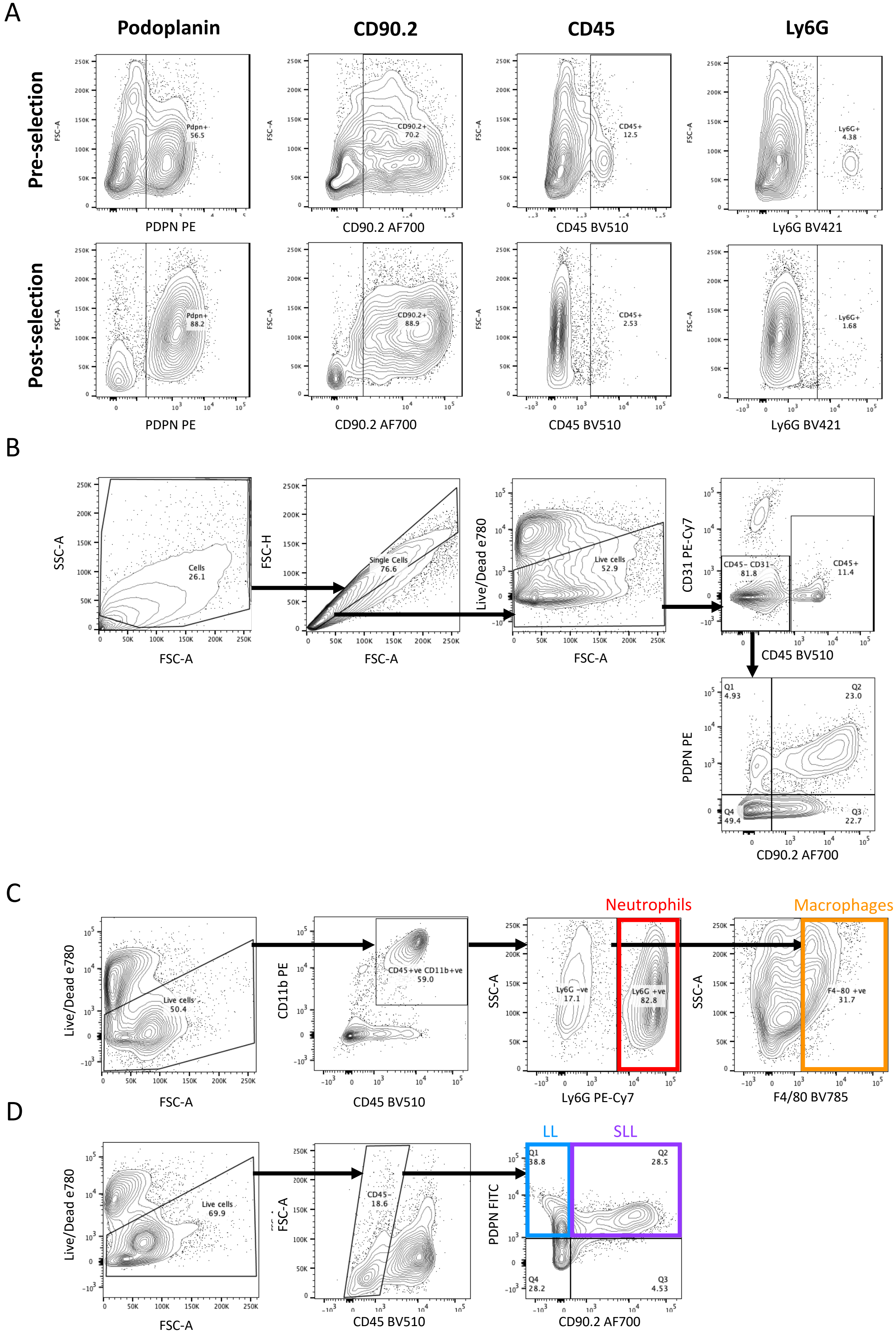


**Supplementary Figure S5.**

**Supplementary Figure S5**

**A.** Flow cytometry analysis of FLS cell enrichment following bead-based purification of $PDPN^+$ cells (bottom), compared to a parallel mock purified sample (top). Plots show single live cell populations.

**B**. Gating strategy for FLS cell sorting. Images are taken from analysis of a representative naïve joint sample. Initial gates for single live cells were applied to all experiments and are omitted from subsequent figures.

**C.** Gating strategy for analysis of CD45+ immune cells in naïve and CIA joint digests. Images are taken from analysis of a representative CIA joint sample. Subpopulations presented in Fig. 3E are highlighted.

**C.** Gating strategy for analysis of Podoplanin+ cells in naïve and CIA joint digests. Images are taken from analysis of a representative CIA joint sample. Subpopulations presented in Fig. 3F are highlighted.

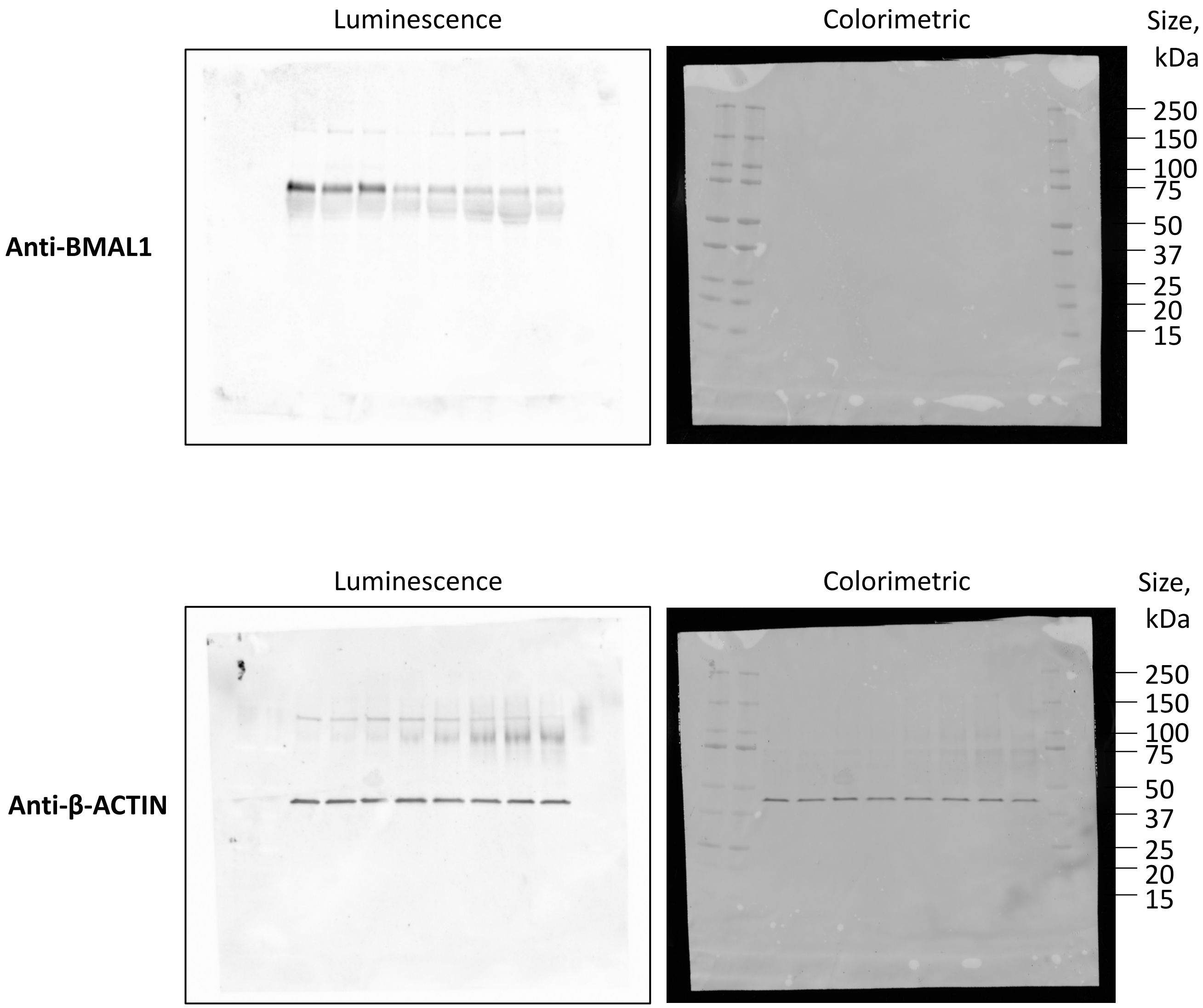


**Supplementary Figure S6.**

**Supplementary Figure S6.**

Uncropped Western blot images.

### Table S1. Primer sequences & qRT-PCR reagents.

| Target | Forward primer (5' to 3') | Reverse primer (5' to 3') | Probe sequence (5' to 3') |
|---|---|---|---|
| Genotyping | | | |
| *Bmal1 flox* | ACTGGAAGTAACTTTATCAAACTG | CTGACCAACTTGCTAACAATTA | |
| *Col6a1 cre* | ATTACCGGTCGATGCAACGAG | CAGGTATCTCTGACCAGAGTC | |
| *Per2::luc* WT | CTGTGTTTACTGCGAGAGT | GGGTCCATGTGATTAGAAAC | |
| *Per2::luc* mut | CTGTGTTTACTGCGAGAGT | TAAAACCGGGAGGTAGATGAGA | |
| *CAG-EYFP* WT | AAGGGAGCTGCAGTGGAGTA | CCGAAAATCTGTGGGAAGTC | |
| *CAG-EYFP* mut | ACATGGTCCTGCTGGAGTTC | GGCATTAAAGCAGCGTATCC | |
| Sybr qRT-PCR | | | |
| *Bmal1* | CCAAGAAAGTATGGACACAGACAAA | GCATTCTTGATCCTTCCTTGGT | |
| *Cry1* | TCGCCGGCTCTTCCAA | TCAAGACACTGAAGCAAAAATCG | |
| *Il1b* | AACCTGCTGGTGTGTGACGTTC | CAGCACGAGGCTTTTTTGTTGT | |
| *Il6* | CCGGAGAGGAGACTTCACAGA | AGAATTGCCATTGCACAACTCTT | |
| *Mmp3* | CTCCTCCACAGACTTGTCCC | AGGACATCAGGGGATGCTGT | |
| *Nr1d1* | GTCTCTCCGTTGGCATGTCT | CCAAGTTCATGGCGCTCT | |
| *Per2* | GCCTTCAGACTCATGATGACAGA | TTTGTGTGCCTCAGCTTTGG | |
| *Ppia* | TATCTGCACTGCCAAGACTGAGTG | CTTCTTGCTGGTCTTGCCATTCC | |
| *Prg4* | CGCCTTTTCCAAAGATCAATACTA | GTGGTAATTGCTCTTGCTGTT | |
| *Tbp* | AGAACAATCCAGACTAGCAGCA | GGGAACTTCACATCACAGCTC | |
| *Thy1* | CACCCCTGGTGAAAACTGCG | AGTTCCGACTTGGATTCTGGAC | |
| Primer/Probe qRT-PCR | | | |
| *Bmal1 (exon 8)* | CGTCGGGACAAAATGAACAG | GAACAGCCATCCTTAGCAC | TACCAACATGCAATGCAATGTCCAGGAA |
| *Actb* | AGGTCATCACTATTGGCAACGA | CACTTCATGATGGAATTGAATGTAGTT | TGCCACAGGATTCCATACCCAAGAAGG |
| *Cxcl1* | CTGCACCCAAACCGAAGTC | AGCTTCAGGGTCAAGGCAAG | CACTCAAGAATGGTCGCGAGGC |
| qRT-PCR with TaqMan gene expression assays | | | |
| *Cre* | | | Mr00635245_cn |

**Table S2. Antibodies for flow cytometry and cell sorting.**

| | Marker | Fluorophore | Channel | Source |
|---|---|---|---|---|
| Sorted FLS | ***Extracellular*** | | | |
| | CD90.2 | AF700 | R 730/45 | BioLegend 105319 |
| | CD45 | BV510 | V 525/50 | BioLegend 103137 |
| | PDPN | PE | Y 586/15 | BioLegend 127407 |
| | CD31 | PE Cy7 | Y 780/60 | BioLegend 102418 |
| Flow analysis of Cre-reporter EYFP joints | ***Extracellular*** | | | |
| | Ly6C | PerCP 5.5 | B 710/50 | BD 560525 |
| | CD90.2 | AF700 | R 730/45 | BioLegend 105319 |
| | Ly6G | BV421 | V 450/50 | BioLegend 127628 |
| | CD45 | BV510 | V 525/50 | BioLegend 103137 |
| | F4/80 | BV785 | V 780/60 | BioLegend 123141 |
| | PDPN | PE | Y 586/15 | BioLegend 127407 |
| | CD31 | PE Cy7 | Y 780/60 | BioLegend 102418 |
| Flow analysis of joint CD45+ immune cells | ***Extracellular*** | | | |
| | CD206 | Per CP 5.5 | B 710/50 | BioLegend 141715 |
| | CD45 | BV510 | V 525/50 | BioLegend 103137 |
| | F4/80 | BV785 | V 780/60 | BioLegend 123141 |
| | CD11b | PE | Y 586/15 | BioLegend 101207 |
| | Ly6G | PE Cy7 | Y 780/60 | BioLegend 127617 |
| | ***Intracellular*** | | | |
| | MerTK | BV421 | V 450/50 | BioLegend 151510 |
| Flow analysis of joint FLS | ***Extracellular*** | | | |
| | PDPN | FITC | B 530/30 | BioLegend 127415 |
| | CD106 | Per CP 5.5 | B 710/50 | BioLegend 105715 |
| | CD55 | APC | R 670/14 | BioLegend 131811 |
| | CD90.2 | AF700 | R 730/45 | BioLegend 105319 |
| | CD45 | BV510 | V 525/50 | BioLegend 103137 |
| | ***Intracellular*** | | | |
| | FAPa | Unconjugated, rat IgG1 | - | R&D MAB9727 |
| | anti-rat IgG | PE | Y 586/15 | R&D F0105B |

### Legends for Supplementary datasets

**Dataset S1. RNAseq analysis of FLS cells isolated from naïve and CIA joints.**

Sheet 1: DEseq2 analysis of differential gene expression.

Sheet 2: EnrichR gene enrichment analysis of top 1000 differentially expressed genes per condition.

**Dataset S2. Single cell RNAseq analysis of FLS cells isolated from *Cre-* and *Cre+* joints.**

Sheet 1: EdgeR analysis of differential gene expression.

Sheet 2: EnrichR gene enrichment analysis of differentially expressed genes.
